# Ocular Safety and Efficacy of AAV-mediated Tyrosinase Gene Augmentation in a Nonhuman Primate Model

**DOI:** 10.64898/2026.07.13.738268

**Authors:** Jaegook Lim, Alessandra M. Larimer-Picciani, Ala Moshiri, Jui-Kai Wang, Noriyoshi Takahashi, Ana Cláudia Santos Raposo, Monica J. Motta, Leah C. Byrne, Sara M. Thomasy

## Abstract

**Purpose:** Oculocutaneous albinism type 1 (OCA1) is an inherited disorder caused by tyrosinase (*TYR*) gene mutations. Affected individuals experience visual impairment and severe photosensitivity from ocular hypomelanosis, with no current treatments. We evaluated the safety and efficacy of a *TYR*-encoding adeno-associated virus (AAV) vector in healthy rhesus macaques as a potential OCA1 treatment.

**Methods:** A novel AAV2-based capsid (ATX002) was packaged with the human VMD2 promoter and *TYR* (h*TYR*) fused with mGreenLantern (mGL). Two adult rhesus macaques were injected with ATX002-hVMD2-h*TYR*-*mGL* subretinally (OD) and intravitreally (OS). Safety and efficacy were assessed via comprehensive ophthalmic examination, fundus photography, spectral-domain optical coherence tomography (SD-OCT), and full-field electroretinography at baseline and defined timepoints up to 12 weeks post-injection, followed by post-mortem immunohistochemistry (IHC).

**Results:** Both subretinal doses induced localized hypermelanosis by 3 weeks post-injection, which persisted through the study endpoint and was accompanied by measurable thickening of the retinal pigment epithelium (RPE) on SD-OCT. Histological IHC confirmed successful RPE transduction via robust mGL fluorescence, corroborating *in vivo* findings by revealing localized RPE hyperplasia and transgene-expressing cells adjacent to regions of *de novo* hypermelanosis. Intravitreal delivery did not induce any changes to the RPE. Transient uveitis was observed but successfully managed with anti-inflammatory treatment.

**Conclusions:** Subretinal AAV-*TYR* delivery is a safe and effective approach with the potential to induce RPE pigmentation. These findings support the use of AAV-*TYR* gene therapy for OCA1, demonstrating efficacy and a manageable safety profile in a large-animal model, and provide a critical bridge toward human clinical translation.

## 1. Introduction

Oculocutaneous albinism (OCA) is a group of autosomal recessive pigmentary disorders characterized by reduced or absent melanin synthesis, resulting in hypopigmentation of the skin, hair, and eyes. The global prevalence of non-syndromic albinism is approximately 1 in 17,000.^1^ Affected individuals experience marked visual dysfunction, including nystagmus, photophobia, and markedly reduced visual acuity, stemming from abnormal retinal development.^2^

Melanin in the retinal pigment epithelium (RPE) is essential for ocular development and light tolerance.^3^ The most common and severe subtype, OCA1, results from biallelic mutations in the tyrosinase gene (*TYR*),^4, 5^ which encodes the rate-limiting enzyme for melanin synthesis in melanocytes and the RPE. Loss of TYR activity disrupts RPE maturation, retinal lamination, and optic nerve fiber decussation, leading to foveal hypoplasia, optic chiasm misrouting, and consequent visual impairment and photosensitivity in OCA1.^6–8^

Currently, management of OCA1-related visual dysfunction focuses on low-vision aides, addressing associated clinical features such as nystagmus, and minimizing the risk of skin-related complications, including skin cancer.^9^ There are currently no curative therapies to reverse OCA1-associated photosensitivity and low visual acuity, making the development of ocular OCA1 therapies a critical and currently unmet medical need.

We postulate that restoration of ocular melanin will ameliorate hypopigmentation-related visual effects, such as photosensitivity, glare, and reduced contrast sensitivity, thereby improving gross visual function and overall quality of life for individuals with OCA1. Adeno-associated virus (AAV)-mediated *Tyr* gene replacement has been evaluated for its potential to restore ocular melanin and improve vision in an OCA1 mouse model.^10, 11^ Recently, we demonstrated that a novel AAV-*Tyr* delivery approach successfully increased ocular pigmentation and reduced photophobic behavior in a well-studied OCA1 murine model.^12^ However, mice lack a fovea and differ anatomically and immunologically from primates. Therefore, evaluation of AAV vectors in a foveated, large-eye model is required for clinical translation, and nonhuman primates (NHPs) provide the ideal translational context. Numerous AAV vectors have been extensively studied in the NHP retina, demonstrating efficient and stable transgene expression with a favorable safety profile prior to clinical use.^13–15^ Additionally, multiple studies have reported spontaneous non-syndromic albinism mutations, including OCA1, in rhesus macaques and other NHP species,^16–18^ which may serve as disease models for future therapeutic studies.

The purpose of this study was to evaluate the transduction efficiency, safety, and biocompatibility of a novel AAV2-based vector, ATX002, derived from our AAV capsid engineering platform^19, 20^, which expresses codon-optimized human *TYR* in the eyes of healthy adult NHPs. Here, we have established a foundational safety and efficacy profile to guide future therapeutic studies in an OCA1 NHP model and support AAV vector clinical translation. Each macaque received a subretinal injection in one eye and an intravitreal injection in the contralateral eye. We hypothesized that intraocular AAV-*TYR* would increase TYR expression in the RPE, with subretinal dosing yielding greater, localized expression compared to intravitreal delivery.

## 2. Materials & Methods

### 2.1 Animals

Two adult rhesus macaques (*Macaca mulatta*) were used in this study at the National Biomedical Research Institute (NBRI). All macaques were born and maintained at the NBRI and determined to be healthy based on ophthalmic examinations, physical assessments, complete blood counts, and serum chemistry profiles. Furthermore, genotyping confirmed that both subjects were wildtype for the *OPA1* (p.A8S) and *PDE6C* (p.R565Q) variants previously identified in this colony.^21, 22^ The NBRI is accredited by the Association for Assessment and Accreditation of Laboratory Animal Care (AAALAC) International.

All procedures adhered to the guidelines of the Association for Research in Vision and Ophthalmology (ARVO) Statement for the Use of Animals in Ophthalmic and Vision Research and the National Institutes of Health (NIH) Guide for the Care and Use of Laboratory Animals. The study protocols were approved by the Institutional Animal Care and Use Committee (IACUC) at the University of California, Davis. Macaques were sedated for ophthalmic examinations and imaging sessions and anesthetized with inhaled isoflurane (Piramal Critical Care, Bethlehem, PA, USA) for vector administration. All procedures were performed under continuous clinical observation and physiological monitoring by a NBRI veterinarian and trained technician.

### 2.2 AAV Vector

Eight macaques were prescreened for anti-AAV2 neutralizing antibodies, which served as a proxy for prior AAV2 exposure, and were enrolled only if titers were <1:10. A newly engineered AAV2-based capsid (ATX002) derived from our AAV engineering platform^19, 20^ was selected for transgene delivery based on its superior infectivity within retinal tissue, including the RPE, compared to AAV2, AAV2.7m8, and similar capsids. The human *TYR* coding sequence was codon-optimized for translation to rhesus macaque and fused to a fluorescent reporter, *mGreenLantern* (mGL), at the C-terminus. The human *VMD2* (*Best1*) promoter was selected to drive transgene expression for its efficiency in RPE.^23^ The NHP-grade ATX002-hVMD2-h*TYR*-*mGL* was purchased from Packgene (Houston, Texas, USA). Prior to packing, all plasmids were sequence-verified through whole plasmid sequencing (Plasmidsaurus, Eugene, OR, USA) and ITR integrity was assessed via restriction digest. Purified AAVs were diluted in sterile buffer containing sodium chloride, sodium phosphate, Poloxamer 188 (0.001%), pH7.3 and were titered using a quantitative PCR method with ITR-binding primers (abm; Richmond, BC, Canada). Sterility testing confirmed the absence of bacterial and fungal contamination, and endotoxin concentrations were <2.00 EU/mL in all preparations.

### 2.3 Injection Procedures

Macaques were pretreated with intramuscular methylprednisolone acetate (Depo-Medrol; Zoetis, Parsippany, NJ, USA; 22.4 mg/kg) administered once weekly and oral cyclosporine (Apotex Corp., Weston, FL, USA; 9 mg/kg) daily, beginning 10 days before injection. All surgical procedures were performed by a single vitreoretinal surgeon (AM) under general anesthesia, and each eye received a different route of administration.

For the right eyes (subretinal injection), two 25-gauge ports (Alcon, Fort Worth, TX, USA) were placed approximately 3 mm posterior to the corneal limbus at the 10 and 2 o’clock meridians. Fundus visualization was achieved using a Hassan-Tornambe Super View disposable contact lens (Oculus Surgical, Port St. Lucie, FL, USA) with a viscoelastic coupling fluid. A bimanual technique was utilized wherein a fiber optic light pipe was inserted via the left port for illumination, and the subretinal injection cannula was inserted via the right port. The cannula tip was advanced to the temporal parafoveal region to create a localized posterior pole bleb. Macaque 1 (10-year-old male) received a low-dose (6×10^10^ vg/eye in 300 µL), and Macaque 2 (9-year-old female) received a high-dose (2×10^11^ vg/eye in 300 µL) using a 25-gauge cannula with a 41-gauge flexible tip (Peregrine Surgical, New Britain, PA, USA). For the left eyes (intravitreal injection), both macaques received an intravitreal dose of 1×10^12^ vg/eye (300 µL), which was set higher than either subretinal dose to enhance the probability of RPE transduction from the vitreous. Intravitreal delivery was performed using a 25-gauge cannula with a 38-gauge flexible tip (MedOne Surgical, Sarasota, FL, USA) through the pars plana, avoiding the lens and retina.

At the conclusion of the procedure, both eyes received subconjunctival injections of ceftazidime (WG Critical Care, Paramus, NJ, USA; 20 mg) and triamcinolone acetonide (Teva Pharmaceuticals, Parsippany, NJ, USA; 8 mg), followed by the topical application of 0.5% erythromycin ophthalmic ointment (Armas Pharmaceuticals, Manalapan, NJ, USA). Post-operatively, macaques received a single dose of sustained-release buprenorphine (Simbadol; Zoetis, Parsippany, NJ, USA; 0.72 mg/kg) and enrofloxacin (Baytril 100; Elanco, Greenfield, IN, USA; 5 mg/kg) twice daily for 5 days. Oral cyclosporine was continued daily for 14 days post-injection, and weekly methylprednisolone acetate injections were continued throughout the study duration. At the 1-week follow up, Macaque 2 received an additional intravitreal injection of dexamethasone (Hikma Pharmaceuticals, Eatontown, NJ, USA; 0.5 mg) in both eyes to address uveitis.

### 2.4 Post-Operative Monitoring

Longitudinal clinical monitoring was performed under sedation with intramuscular ketamine (Zetamine; Vet One, Boise, ID, USA; 10 mg/kg), dexmedetomidine (Dexmedesed; Dechra, Overland Park, KS, USA; 0.03 mg/kg), and midazolam (Hikma, Berkeley Heights, NJ, USA; 0.1 mg/kg), supplemented with additional ketamine and dexmedetomidine as needed to maintain adequate sedation.

Comprehensive ophthalmic examinations and retinal imaging were conducted at baseline, and at 1, 3, 6, and 12 weeks post-injection to assess transgene expression, melanin production, ocular inflammation, and signs of toxicity. Ocular safety and toxicity were quantified using the semiquantitative preclinical ocular toxicology scoring (SPOTS) system.^24^ Full-field electroretinography (ERG) was performed at baseline, 3, 6, and 12 weeks post-injection to evaluate retinal function.

Intraocular pressure (IOP) was measured at all time points using rebound tonometry (TonoVet; Icare, Helsinki, Finland).^25^ Measurements were obtained with the macaques gently restrained in an upright position, and only readings with minimal deviation were accepted in accordance with the manufacturer’s recommendations. Following IOP measurement, pupils were dilated with 1% tropicamide (Bausch & Lomb, Tampa, FL, USA) and 2.5% phenylephrine (Akorn, Lake Forest, IL, USA). Slit-lamp biomicroscopy (SL-17; Kowa, Tokyo, Japan) and indirect ophthalmoscopy (Vantage Plus; Keeler, Broomall, PA, USA) with a 20 or 28D indirect lens (Volk Optical; Mentor, OH, USA) were then performed. At the conclusion of each session, corneal integrity was assessed with fluorescein staining (BioGlo; HUB Pharmaceuticals, Rancho Cucamonga, CA, USA).

### 2.5 Optical Coherence Tomography and Retinal Imaging

Retinal imaging was performed following previously established methods.^25–27^ At each examination, color fundus photographs were obtained using a 50° wide-angle retinal camera (CF-1; Canon, Tokyo, Japan) equipped with a digital camera (EOS 5D; Canon, Tokyo, Japan). Spectral-domain optical coherence tomography (SD-OCT), blue-light autofluorescence (BAF), and fluorescein angiography were conducted using a confocal scanning laser ophthalmoscope (Spectralis HRA+OCT; Heidelberg Engineering, Heidelberg, Germany). An eyelid speculum was inserted to facilitate imaging, and corneal hydration was maintained with topical artificial tears (Genteal; Alcon, Fort Worth, TX, USA) at approximately 2-minute intervals or as required.

Quantitative analysis of retinal layer thickness was performed on SD-OCT images using Heidelberg Eye Explorer software (Heidelberg Engineering). The foveal center was identified as the point of maximum retinal depression, and ImageJ software (version 1.54i; National Institutes of Health, Bethesda, MD, USA) was used to calibrate the images based on the manufacturer’s scale prior to manual measurement. Retinal layer thicknesses were measured at the foveal center and at 1.5 mm temporal and nasal to the fovea. The following layers were analyzed: retinal nerve fiber layer (RNFL), ganglion cell layer (GCL), inner plexiform layer (IPL), inner nuclear layer (INL), outer plexiform layer (OPL), outer nuclear layer (ONL), photoreceptor inner segments (IS), photoreceptor outer segments (OS), RPE, choriocapillaris (CC), outer choroid (OC), and total retinal thickness (TRT).^25, 27^

### 2.6 Electroretinography (ERG)

The ERG recordings were obtained using a handheld system (Retevet; LKC Technologies, Gaithersburg, MD, USA). Macaques were dark-adapted for 30 minutes under sedation prior to testing. Flash stimuli of 0.01, 3.0, and 10.0 cd·s/m² were presented in the dark-adapted state, followed by light adaptation for 10 minutes. Photopic testing included flash stimuli at 3.0 cd·s/m² and a 30-Hz flicker response. Photopic negative responses (PhNR) were recorded using a red flash (1.0 cd·s/m² at 3.4 Hz) on a blue background (10 cd·s/m²). All recordings were analyzed using the manufacturer’s software to determine implicit time (ms) and amplitude (μV) for each eye across all testing conditions.

### 2.7. Tissue Processing and *Ex Vivo* Analysis

Serum was collected at baseline, 1, 6, and 12 weeks post-injection to evaluate the systemic immune response to the viral vector used through the quantification of anti-AAV2 neutralizing antibody (NAb) titers. Serum was sent to VRL Animal Health Diagnostics (San Antonio, Texas, USA) for titer quantification. Titers were quantified as the greatest sera dilution capable of inhibiting ≥ 50% AAV2 transduction *in vitro*. At 12 weeks post-injection, macaques were euthanized, and ocular tissues were collected for histological analyses. Whole eyes were fixed in 4% paraformaldehyde for 18 hours, transferred to PBS, and stored at 4□ thereafter. Post-fixation, the anterior segment was removed and a manual dissection of the vitreous was performed at the ora serrata. Eye cups were cryoprotected in 15% and 30% sucrose/DI H_2_O solutions. Prior to embedding in Optimal Cutting Temperature compound (Fisher Scientific, Hampton, NH, USA), cryoprotected eye cups were divided into temporal and nasal segments via a vertical incision at the optic nerve. 16-µm frozen sections were collected and stored at −80□ until further processing. Cryosections were adhered to the slide with 4% PFA for 10 minutes. To reduce background autofluorescence, slides were incubated in 1 mg/mL NaBH_4_ for 10 minutes before being stained with Hoechst 33342 (1:2000; Thermo Fisher Scientific, Waltham, MA, USA) nuclear stain. Slides were mounted with Fluoromount-G™ Mounting Medium (Invitrogen, Carlsbad, CA, USA) and imaged using an Olympus Fluoview 4000 confocal microscope (Olympus Corporation, Tokyo, Japan). Median intensity Z projections were created in ImageJ.

### 2.8. Data analysis

Due to the small sample size, the results of this study are presented descriptively. Quantitative data from *in vivo* analyses were compared to their respective baseline measurements. The results were also compared with previously collected data from 17 healthy, age-matched rhesus macaques (mean age±standard deviation [SD]: 7.5±1.4 years) from the same NBRI colony.^25^ Graphs were generated using GraphPad Prism (version 10.6.1; GraphPad Software, Boston, MA, USA).

## 3. Results

### 3.1 Subretinal and Intravitreal AAV-*TYR* Delivery is Well Tolerated with a Manageable Safety Profile

While IOP initially decreased in all eyes after injection, it remained within the established reference range and was comparable to those of age-matched rhesus macaques (**Figure 1**).^25, 27^ Specifically, IOP ranged from 9–22 mmHg in subretinally-injected eyes and from 9-24 mmHg in intravitreally-injected eyes.

**Figure 1.**
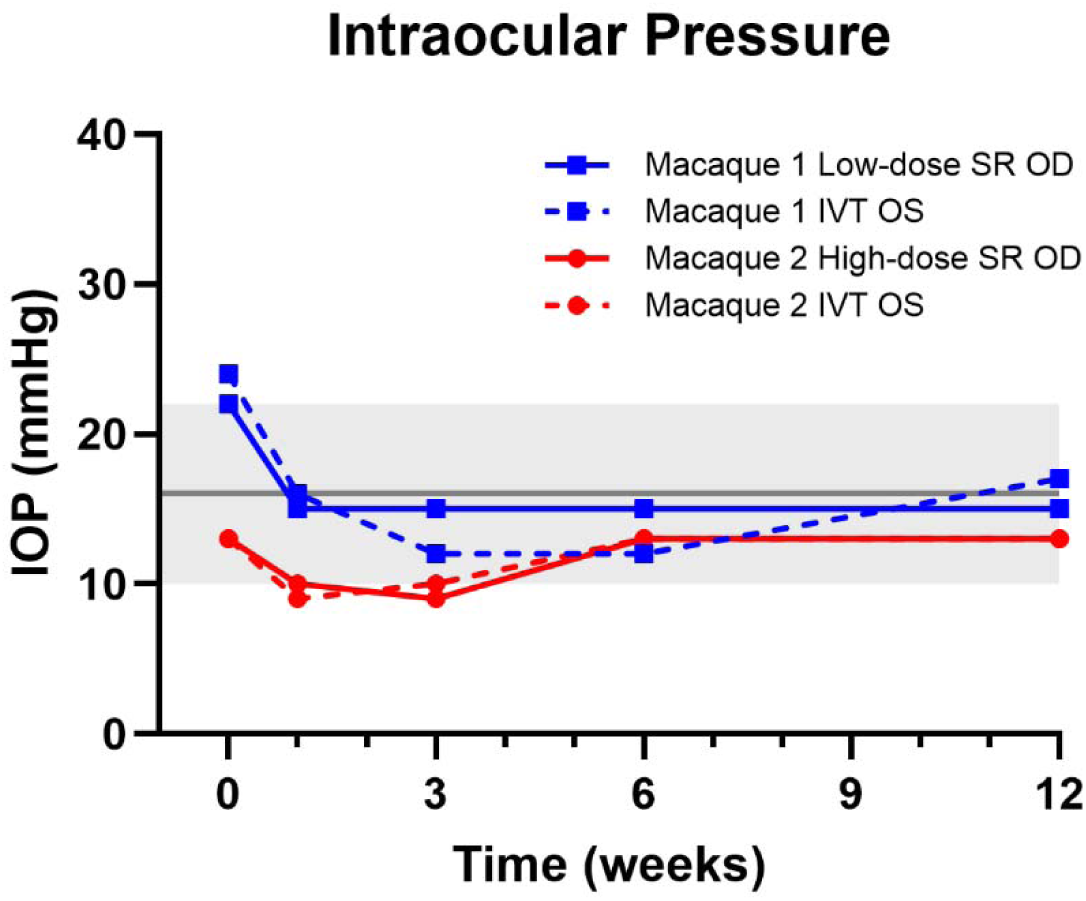
Intraocular pressure (IOP) transiently decreased following subretinal and intravitreal ATX002-hVMD2-h*TYR*-*mGL* injection then remained within the reference range for the remainder of the study period. Blue and red lines represent Macaque 1 and 2, respectively. Measured values are compared with the reference mean (solid dark gray horizontal line) and minimum to maximum range (gray shading) from a previously reported cohort of normal rhesus macaques (mean age ± SD, 7.5 ± 1.4 years; n = 17) from the same colony, obtained using identical equipment and methods.^25^

With SD-OCT imaging, we confirmed preservation of all retinal layers in all eyes throughout the 12-week study period. Specifically, SD-OCT measurements 1.5 mm temporal to the fovea showed a temporary decrease in OS thickness in subretinally-injected eyes from 1 to 3 weeks post-injection, which recovered by 6 weeks (**Figure 2**). We also observed transient attenuation of the ellipsoid zone (EZ) near the fovea in the low-dose subretinally-injected eye at 1-week post-injection, which resolved by 3 weeks post-injection (**Supplementary Figure 1**). The TRT in the subretinally-injected eyes also showed a transient decrease at 1-week post-injection but recovered to baseline by 6 weeks post-injection and maintained through the 12-week endpoint. The ONL thickness remained reduced compared to baseline values through the 12-week timepoint in both eyes (high-dose subretinal and intravitreal) of Macaque 2, as well as the low-dose subretinally injected eye of Macaque 1; however, all values remained within the reference range established from non-injected age-matched rhesus macaques.^25^ All other retinal layers (RNFL, GCL, IPL, INL, and OPL) remained stable, demonstrating minor fluctuations in thickness. Finally, fluorescein angiography images revealed a hyperfluorescent region at the site of high-dose subretinal injection (**Figure 3B 3**).

**Figure 2.**
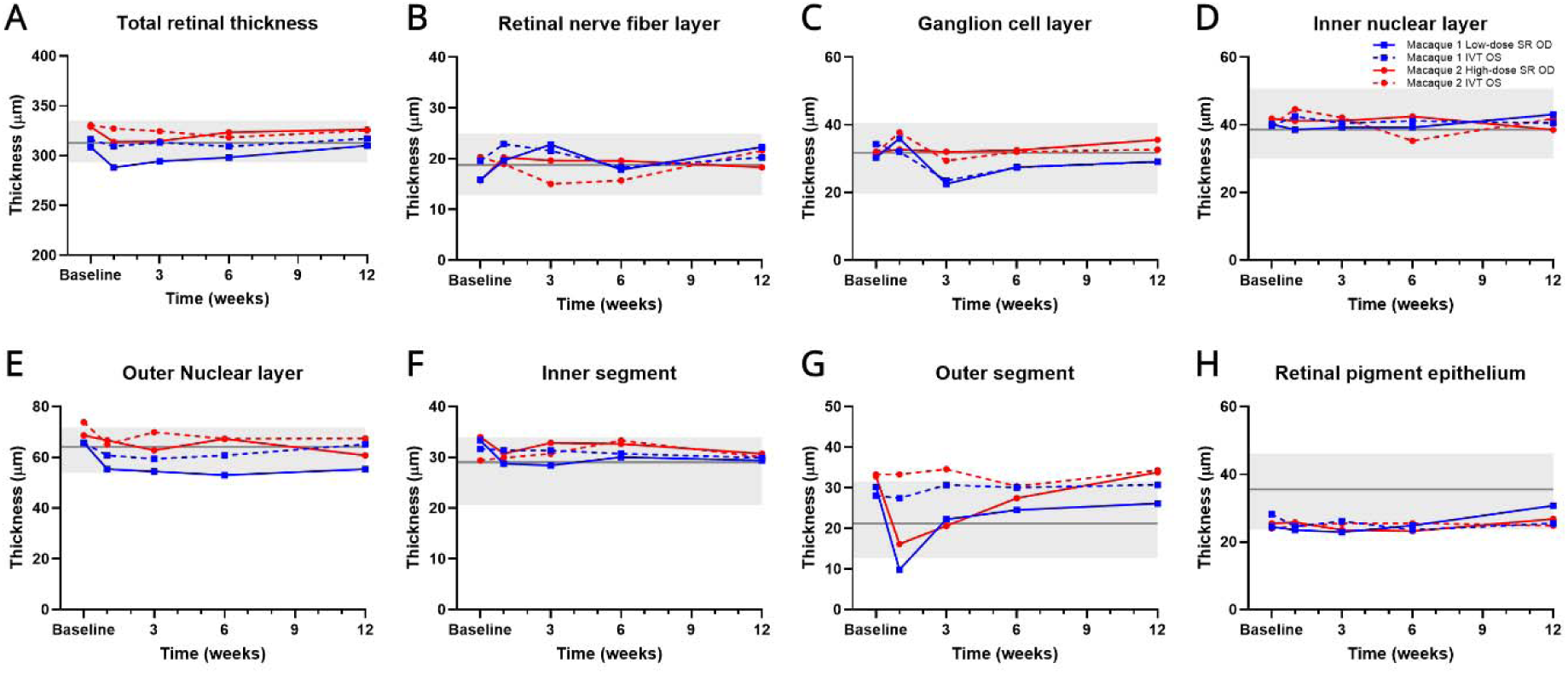
Transient parafoveal outer segment thinning was observed at 1 and 3 weeks following subretinal and intravitreal ATX002-hVMD2-h*TYR*-*mGL* injection, which resolved over time. Retinal layer thickness measured 1.5 mm temporal to the fovea, including (A) total retinal thickness, (B) retinal nerve fiber layer, (C) ganglion cell layer, (D) inner nuclear layer, (E) outer nuclear layer, (F) inner segment, (G) outer segment, and (H) retinal pigment epithelium. Measurements are shown for AAV-*TYR* treated eyes up to 12 weeks post-injection. Blue and red lines represent Macaque 1 and Macaque 2, respectively. Measured values are compared with the reference mean (solid dark gray horizontal line) and minimum to maximum range (gray shading) from a previously reported cohort of normal rhesus macaques (mean age ± SD, 7.5 ± 1.4 years; n = 17) from the same colony, obtained using identical equipment and methods.^25^

**Figure 3.**
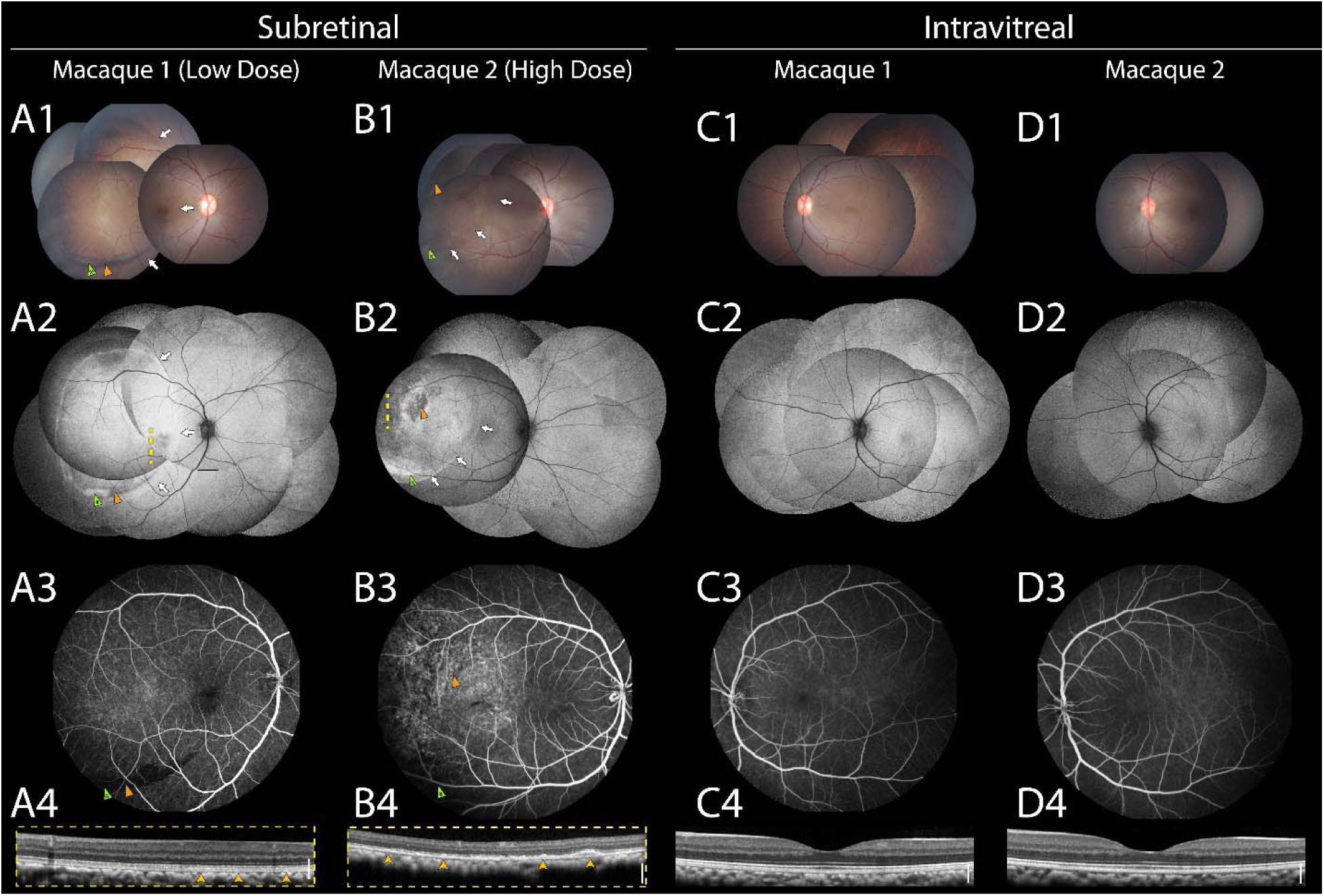
Subretinal delivery appears to induce localized melanogenesis and transgene expression without vascular leakage. Comprehensive multimodal imaging of subretinally (low- and high-dose) and intravitreally injected eyes at 12 weeks post-injection is shown. Columns are grouped by injection method: subretinally injected eyes (Macaque 1 [low-dose], **A**; Macaque 2 [high-dose], **B**) and their respective intravitreally injected fellow eyes (Macaque 1, **C**; Macaque 2, **D**). Rows display color fundus montages (**1**), blue autofluorescence montages (**2**), fluorescein angiography (**3**), and corresponding SD-OCT scans (**4**). In the subretinally injected eyes (**A**, **B**), white arrows indicate the margins of the subretinal bleb. Green arrowheads (marked with asterisks) identify areas of mGreenLantern expression, while orange arrowheads indicate regions of *de novo* melanin synthesis. The yellow dashed lines in **A2** and **B2** indicate the scan locations for the OCT images shown in **A4** and **B4**, respectively, demonstrating retinal pigment epithelium thickening (yellow arrowheads). **C4** and **D4** show representative foveal scans of the intravitreally injected fellow eyes, displaying normal retinal architecture. Scale bars indicate 200 µm.

Mixed rod-cone responses, as well as light-adapted cone and flicker amplitudes and implicit times, demonstrated minor fluctuations over time but generally returned to levels within the range of age-matched healthy macaques by the 12-week endpoint, except for a prolongation of the b-wave implicit time observed at the 12-week endpoint in the low-dose subretinally-injected eye (**Figure 4**).

**Figure 4.**
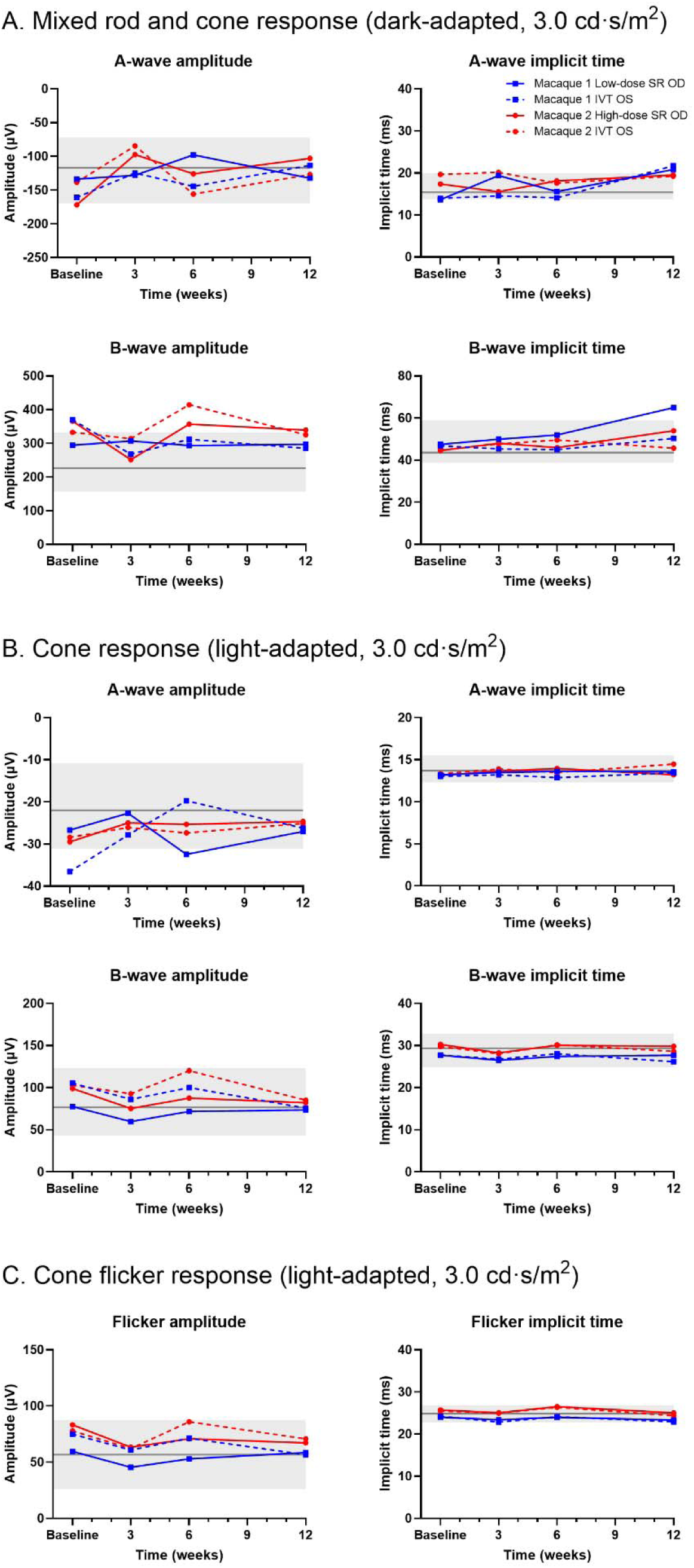
Retinal function was preserved following subretinal and intravitreal ATX002-hVMD2-h*TYR*-*mGL* injection, with electroretinography (ERG) parameters remaining comparable to the normative range. Line plots showing full-field ERG measurements of a- and b-wave amplitudes (left panels) and implicit times (right panels) after dark adaptation using (A) 3.0 cd·s/m² at 0.1 Hz, and after light adaptation using (B) 3.0 cd·s/m² at 2 Hz and (C) at 28.3 Hz. Blue and red lines represent Macaque 1 and 2, respectively. Measured values are compared with the reference mean (solid dark gray horizontal line) and minimum–maximum range (gray shading) from a previously reported cohort of normal rhesus macaques (mean age ± SD, 7.5 ± 1.4 years; n = 17) from the same colony, obtained using identical equipment and methods.^25^

Macaque 2 (high-dose subretinal injection in the right eye) initially developed moderate uveitis in both eyes. At 1-week post-injection, the subretinally-injected eye exhibited dense white cells within the vitreous, mild papilledema, and focal vitreous hemorrhage. This inflammation improved by 6 weeks and completely resolved by 12 weeks post-injection in the subretinally-injected eye, while mild pigmented vitreous cells remained in the intravitreally-injected eye. By contrast, Macaque 1 exhibited no signs of inflammation in the low-dose subretinally-injected eye, whereas the intravitreally-injected eye showed mild inflammation with white and brown vitreous cells observed at 1-week post-injection, which completely resolved by 12 weeks post-injection.

Macaque 2 (high subretinal dose) demonstrated greater peak AAV2 Nab titers at 1 and 6 weeks post-injection compared to Macaque 1 (low subretinal dose), consistent with the aforementioned clinical findings (**Table 1**). AAV2 NAb titer values returned to baseline (<1:10) by the 12-week endpoint.

**Table 1.** AAV2 neutralizing antibody titers return to baseline by 12 weeks post-injection.

| Subject | Vector dose<br>(vg/eye) | Baseline AAV2<br>titer | 1 week post-op<br>AAV2 titer | 6 week post-op<br>AAV2 titer | 12 week post-op AAV2 titer |
| --- | --- | --- | --- | --- | --- |
| **Macaque 1** | 6×10<sup>11</sup> OD<br>1×10<sup>12</sup> OS | <1:10 | 1:93 | 1:10 | <1:10 |
| **Macaque 2** | 2×10<sup>11</sup> OD<br>1×10<sup>12</sup> OS | <1:10 | 1:178 | 1:253 | <1:10 |

### 3.2 Subretinal AAV-*TYR* Delivery Mediates Robust Transgene Expression, *De Novo* Melanin Synthesis, and Associated RPE Structural Changes

At 1-week post-injection, mGL expression was already visible in the high-dose subretinally-injected eye, whereas no fluorescence was detected in the low-dose subretinally-injected eye. Distinct dark-brown pigmentation suggestive of melanin deposition became evident within the subretinal bleb region by 3 weeks post-injection in both subretinally-injected eyes (**Figure 5**). Focal melanin accumulation was evident on fundus examination and OCT imaging, becoming progressively darker and more consolidated by 6 weeks and persisted through the 12 weeks in the low- and high-dose subretinally-injected eyes (**Figures 3 and 5**). The greatest RPE thickness was observed in the high-dose subretinally-injected eye (79.1 µm), which exceeded that of the low-dose subretinally-injected eye (49.0 µm) and was higher than the established mean ± SD RPE thickness from age-matched controls at NBRI (33.2 ± 3.7 µm at the foveal center, 35.5 ± 4.6 µm at 1.5 mm temporal to the fovea, and 35.3 ± 3.6 µm at 1.5 mm nasal to the fovea).^25^ The melanin deposition obscured mGL fluorescence in these regions, resulting in a patchy appearing distribution (**Figure 3B2**). No pigmentary changes were observed in any intravitreally-injected eyes.

**Figure 5.**
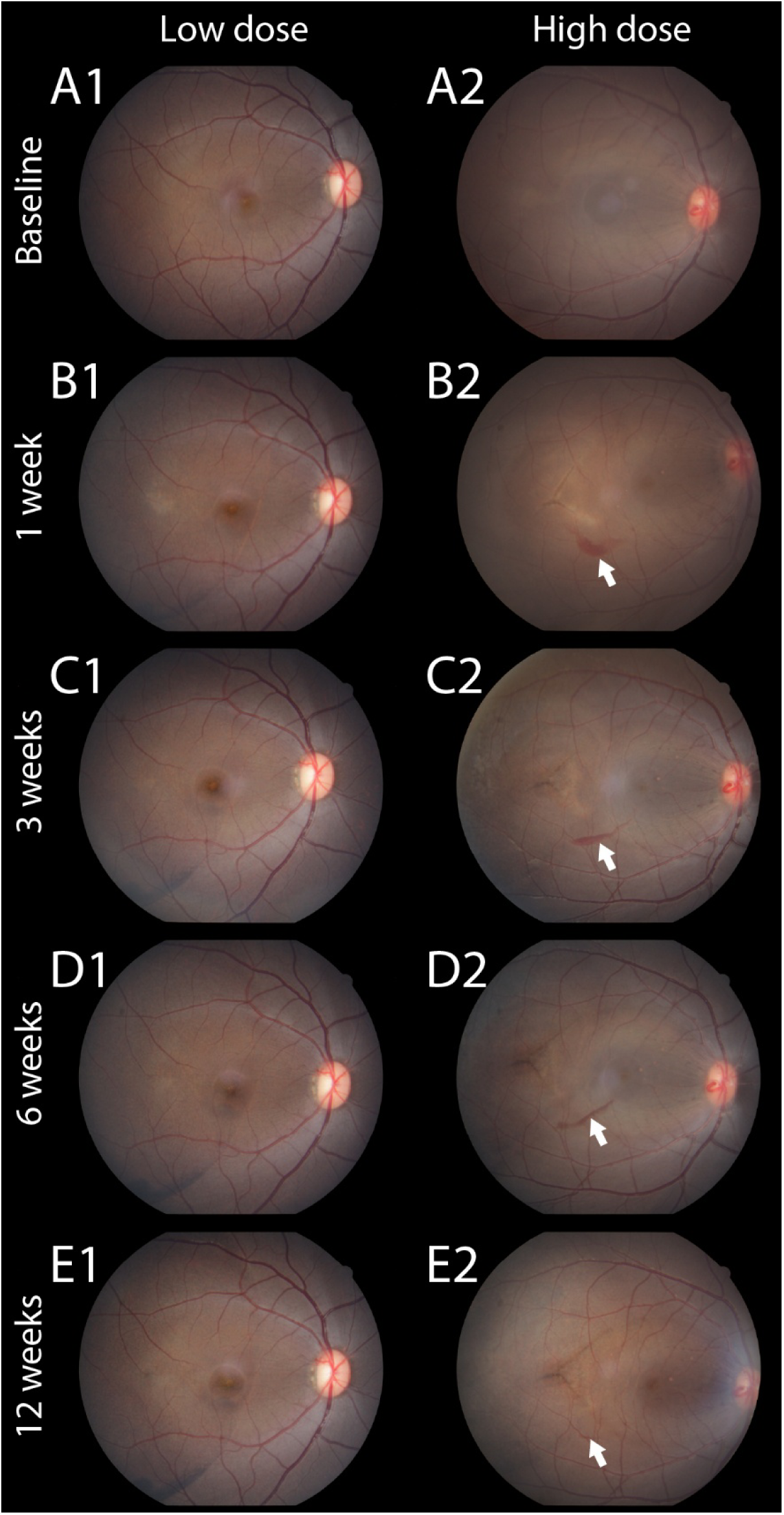
Subretinal ATX002-hVMD2-h*TYR*-*mGL* injection resulted in changes suggestive of progressive localized melanogenesis in the nonhuman primate retina. Longitudinal color fundus photographs of the macula are shown for the low-dose (left column, A1–E1) and high-dose (right column, A2–E2) eyes at the baseline, 1, 3, 6, and 12 weeks post-injection. Dark-brown pigmentation, consistent with melanin, progressively developed, becoming distinct within the subretinal bleb region by 3 weeks (C1 and C2), and persisted through the 12-week endpoint (E1 and E2). White arrows indicate a localized vitreous hemorrhage observed in the high-dose eye starting at 1 week, which markedly diminished by the 12-week endpoint.

At 12 weeks post-injection, subretinally-injected eyes exhibited distinct, localized RPE thickening that formed a consistent spatial pattern in the periphery, as visualized in both horizontal and vertical *en-face* OCT images (**Figure 6**). This regional thickening was reproducible across different scanning orientations and clearly delineated on the montaged RPE thickness map (**Figure 6C2**). The RPE boundaries were obtained by manually refining the segmentation results from an updated deep learning-based approach.^28^

**Figure 6.**
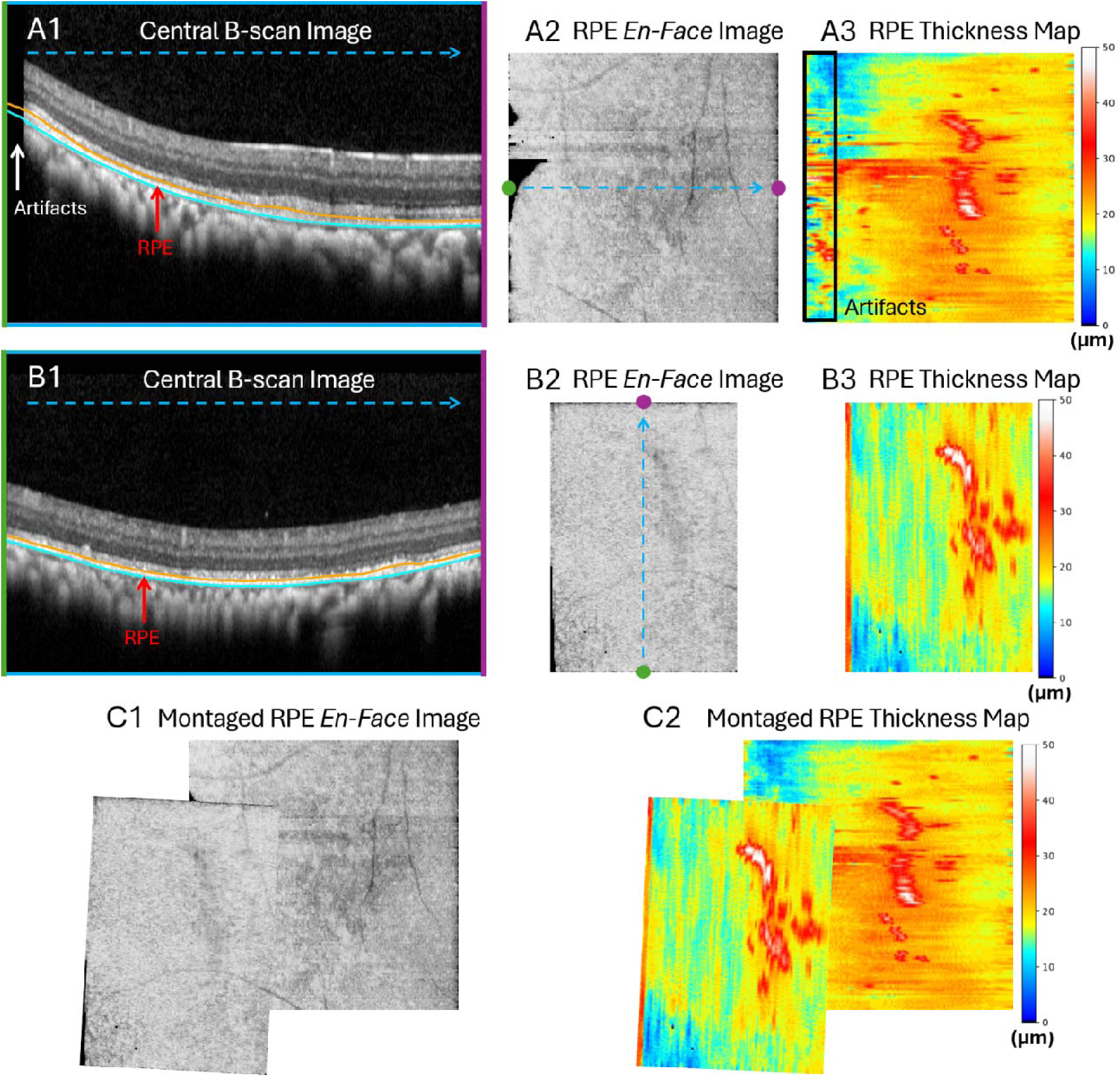
Localized RPE thickening is observed in peripheral retinal regions from two 3D OCT volumes of Macaque 2’s subretinally-injected eye at 12 weeks post-injection. Row. **A** is an OCT volume scanned horizontally covering 4×4 ×2 mm^3^ with 768×144×496 voxels; **Row B** is an OCT volume scanned vertically covering 2.8×4×2 mm^3^ with 100×768×496 voxels. Artifacts are apparent in the OCT volume shown in **Row A**, and the volume in **Row B** wa acquired to provide additional coverage of the affected region. **A1** and **B1** show the central B-scan images of each OCT volume. **A2** and **B2** display the corresponding RPE projection (i.e., *en-face*) image respectively, where the dashed arrows from the green to purple markers indicate the image orientation and location of **A1** and **B1**. **A3** and **B3** display the corresponding RPE thickness maps. **C1** shows the montaged *en-face* image, indicating the spatial relationship between the two OCT volumes, and **C2** displays the corresponding montaged RPE thickness map. Consistent spatial patterns can be observed in both the *en-face* images and the thickness maps.

Confocal fluorescence microscopy of histological sections demonstrated mGL fluorescence in the temporal RPE of rhesus macaques subretinally treated with low- and high-dose ATX002-VDM2-h*TYR*-*mGL*, indicating RPE viral transduction occurred in the regions of interest described above (**Figure 7**). Corroborating the *in vivo* OCT findings of RPE thickening, histological examination also revealed areas of RPE hyperplasia and cellular alterations within these transduced regions (**Figures 7B6**)

**Figure 7.**
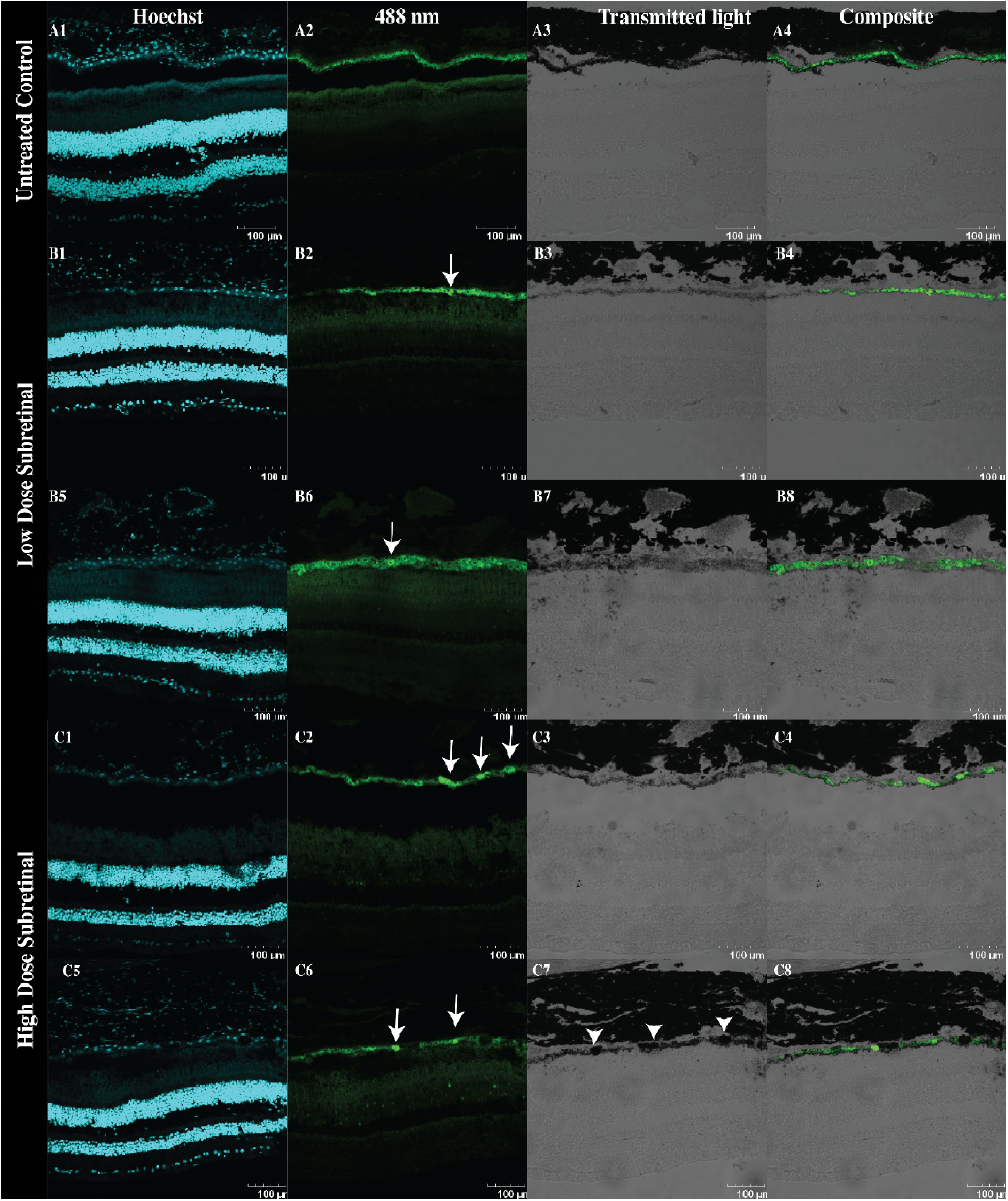
Robust mGreenLantern (mGL) expression and localized retinal pigment epithelium (RPE) hyperplasia in the temporal RPE of rhesus macaques following subretinal ATX002-hVMD2-h*TYR*-*mGL* delivery. A1-A4) Control tissue obtained temporal to the optic nerve from a seven-year-old untreated, control rhesus macaque demonstrated uniform autofluorescence in the RPE at 488 nm (A2). **B1-B8**) The low-dose subretinally-injected eye demonstrated regions of 488 nm signal intensity above background autofluorescence in the RPE in multiple temporal fields of view, indicative of mGL expression (B2, B6). The appearance of RPE hyperplasia was noted in one field of view (B6). **C1-C8)** The high-dose subretinally-injected eye demonstrated localized 488 nm signal intensity above background fluorescence in multiple fields of view (C2, C6), indicating mGL expression in these RPE. Certain fluorescent cells were noted to be adjacent to regions of hypermelanosis (C7). Arrows indicate regions of 488 nm signal intensity and mGL expression. Arrowheads indicate regions of hypermelanosis. All images were acquired at 20X magnification. Scale bars indicate 100 µm.

## 4. Discussion

This study is an essential first step to developing gene therapy for OCA1. A single subretinal injection of the AAV-*TYR* vector induced changes suggestive of melanin synthesis in the RPE of adult macaques, providing key proof-of-concept evidence for an AAV-*TYR*-mediated pigment induction strategy in a large-animal model with ocular anatomy and physiology similar to humans. Subretinal delivery demonstrated superior efficacy to intravitreal administration, as no RPE pigmentation was observed following intravitreal injection. The observations described here support the use of subretinal delivery over intravitreal administration for efficient RPE transduction, given the restrictive nature of inner retinal barriers.^29, 30^

The use of an NHP model was essential for the translational relevance of this study. Unlike mice and other non-foveated preclinical models, the NHP retina is structurally, functionally, and immunologically similar to that of humans, making it the preferred model for retinal AAV translational studies. Importantly, NHPs possess a fovea, the anatomical structure responsible for high-acuity vision.^31^ Because foveal hypoplasia is a defining and debilitating feature of OCA1, the NHP model uniquely enables both safety and efficacy assessments that are directly relevant to human clinical translation in albinism. Crucially, our longitudinal SD-OCT analysis confirmed that foveal architecture was preserved following vector administration. Even in instances where the subretinal bleb directly involved the fovea (specifically in the low-dose injected eye), quantitative analysis demonstrated that the foveal thickness at the 12-week endpoint returned to baseline and remained within the established reference range derived from age-matched controls in the same colony using identical imaging protocols.^25^ These findings support the safety of subretinal AAV delivery in the primate macula and its potential for human clinical translation in albinism.

Findings consistent with localized RPE hypermelanosis and a corresponding increase in RPE thickness became visible at 3 weeks post-injection and persisted through the 12-week endpoint in subretinally-injected eyes. The RPE pigment changes observed in this study were non-uniform, with discrete hyperreflective regions and localized thickening distributed throughout the subretinal bleb region in low- and high-dose eyes. Similarly, confocal fluorescence microscopy revealed mGL positive RPE cells temporal to the optic nerve. A region of RPE hyperplasia was noted in one field of view in the low-dose eye, potentially contributing to the RPE thickness measurements *in vivo* in this eye. The high-dose eye exhibited discrete regions of RPE hypermelanosis, which correlate to the non-uniform increase in RPE thickness observed on OCT in this eye. The patchy appearance of suspected vector-mediated pigment and mGL was likely attributable to both pre-existing pigmentation and variable AAV transduction across the RPE. The macaques selected for this study were healthy rhesus macaques with fully pigmented iris, RPE, and choroidal tissues. We postulate that a high vector genome dose per cell was required to produce sufficient exogenous TYR to generate the hyperpigmentation and increased thickness observed in select RPE regions. These findings may demonstrate a dose-dependent effect, where the higher subretinal dose induced more robust, extensive melanin synthesis accompanied by greater RPE thickening compared to the lower dose. Nonetheless, the presence of mGL-positive RPE cells in subretinally-injected eyes on confocal fluorescence microscopy indicates that ATX002-hVMD2-h*TYR*-*mGL* transduced primate RPE, suggesting melanin changes were related to exogenous TYR activity and not procedure-related changes.

A transient decrease in the thickness of the OS layer was observed in both low- and high-dose subretinally-injected eyes. The thinning observed between 1 and 3 weeks post-injection appears to be a reversible surgical effect, consistent with a previous report in NHPs.^32^ This acute structural change is likely a temporary consequence induced by the surgical bleb. A prior subretinal injection study in NHPs demonstrated that although subretinal delivery induces a transient retinal bleb, the foveal and outer retinal architecture subsequent to vector administration undergoes complete structural recovery without clinically significant permanent thinning, supporting the transient nature of this change.^32^ It has been suggested that these findings represent clinical signs of reversible photoreceptor outer segment alteration associated with the surgical fluid bleb, causing acute but reversible photoreceptor stress. Consistent with these findings, we also observed transient attenuation of the EZ near the fovea in the low-dose subretinally-injected eye at 1-week post-injection, which subsequently recovered. The transient, reversible thinning observed in our study further supports successful and localized subretinal delivery, confirming accurate bleb formation at the intended site.

The hyperfluorescent area observed within the subretinal injection site of the macaque receiving the high-dose suggests a localized alteration of the outer retinal integrity. Notably, an intravitreal injection of steroids was necessary to control inflammation in the high-dose macaque (Macaque 2), whereas only mild inflammation was observed in the low-dose macaque (Macaque 1). Given that this phenomenon was undetected following the low-dose subretinal injection, these findings may reveal a potential dose-dependent risk of localized outer retinal changes. Although these manifestations warrant careful consideration regarding safety, further research focusing on varied concentrations and delivery volumes is warranted to better define the safety profile and optimize the therapeutic window of this gene therapy configuration.

This pilot study has several limitations. First, the small sample size prevents robust statistical analysis and limits the generalizability of the findings. Future studies could address this by increasing the number of macaques and using a paired-eye design, in which one eye receives treatment and the contralateral eye serves as an internal control, enabling more rigorous within-subject comparisons and statistical analysis. Second, the 12-week follow-up period, although sufficient to assess initial safety and biological activity, may not capture potential late-onset toxicities or long-term stability of transgene expression. Finally, the use of healthy macaques precludes evaluation of therapeutic efficacy in correcting the visual deficits characteristic of albinism. Recently, rhesus macaques carrying variants in melanogenesis-related genes, including *TYR*, *OCA2*, and *TYRP1*, have been identified.^18, 33^ Although these macaques present with ‘albino’ or ‘golden’ phenotypes, they typically exhibit residual melanin synthesis characterized by reddish or golden pelage, distinguishing them from the complete loss of pigmentation observed in severe human OCA1. While a spontaneous NHP model with foveal hypoplasia has been reported,^18^ a model with functionally characterized visual compromise remains to be fully established. The application of large-scale genomic resources, such as the Macaque Genotype and Phenotype (mGAP) database,^33, 34^ will be essential to screen for pathogenic *TYR* variants and identify appropriate models of OCA1 for future efficacy studies.

In conclusion, this pilot study provides foundational preclinical evidence of both safety and biological activity for subretinal AAV-*TYR* gene therapy in NHPs. Subretinal administration of AAV-*TYR* safely resulted in changes suggestive of melanogenesis in the RPE of NHPs throughout the 12-week period, establishing a manageable preliminary safety profile in the NHP eye. These results justify advancing this approach into a genetically and phenotypically confirmed NHP model of albinism to further evaluate its therapeutic potential for visual restoration and its translational relevance to patients with OCA1.

## Acknowledgments

We thank the veterinary and technical staff at the NBRI for their invaluable assistance. The authors would like to express their sincere gratitude to Dr. Glenn Yiu for his invaluable expertise and assistance in the interpretation of retinal images. We are grateful to Dr. Chrisoula Toupadakis Skouritakis for her exceptional support in figure preparation and design.

