## Supplementary figures and images for "Ocular Safety and Efficacy of AAV-mediated Tyrosinase Gene Augmentation in a Nonhuman Primate Model"

### Supplementary Figure 1

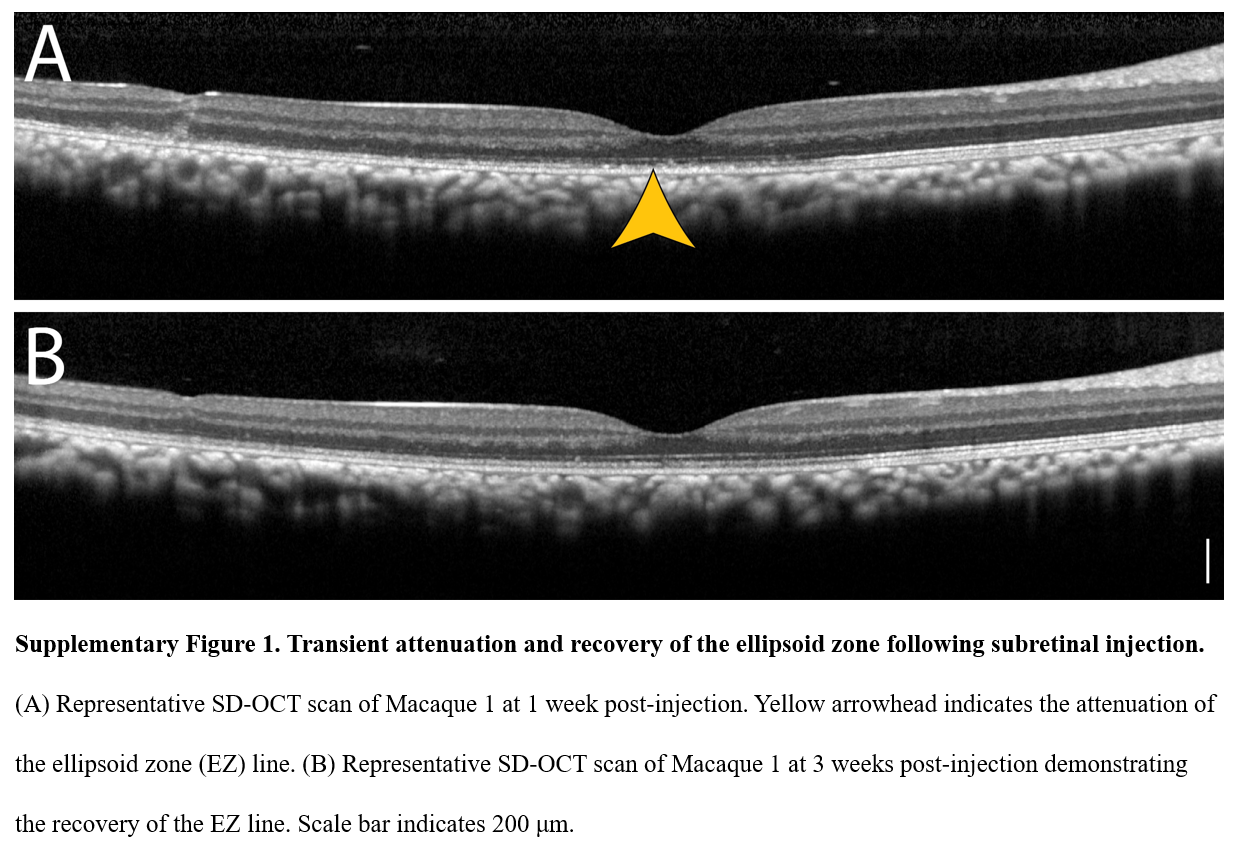
