## Supplementary Table 1 for "Ocular Safety and Efficacy of AAV-mediated Tyrosinase Gene Augmentation in a Nonhuman Primate Model"

**Supplementary Table 1. Summary of longitudinal clinical ophthalmic examination findings following intraocular ATX002-hVMD2-h*TYR*-*mGL* injection.**

| **Subject** | **Route** | **Parameter Scored** | **Baseline** | **1 Week** | **3 Weeks** | **6 Weeks** | **12 Weeks** |
| --- | --- | --- | --- | --- | --- | --- | --- |
| **Macaque 1** | **Subretinal** | Anterior Chamber Flare | 0 | 0 | 0 | 0 | 0 |
|  |  | Anterior Chamber Cell | 0 | 0 | 0 | 0 | 0 |
|  |  | Anterior Vitreous Cell | 0 | 0 | 0 | 0.5 (White) | 0 |
|  |  | Degraded Fundus View | 0 | 0 | 0 | 0 | 0 |
|  | **Intravitreal** | Anterior Chamber Flare | 0 | 0 | 0 | 0 | 0 |
|  |  | Anterior Chamber Cell | 0 | 0 | 0 | 0 | 0 |
|  |  | Anterior Vitreous Cell | 0 | 1 (Mixed) | 1 (White) | 1 (White) | 0 |
|  |  | Degraded Fundus View | 0 | 0 | 0 | 0 | 0 |
| **Macaque 2** | **Subretinal** | Anterior Chamber Flare | 0 | 0 | 0 | 0 | 0 |
|  |  | Anterior Chamber Cell | 0 | 0 | 0 | 0 | 0 |
|  |  | Anterior Vitreous Cell | 0 | 3 (Mixed) | 3 (Mixed) | 3 (Brown) | 0 |
|  |  | Degraded Fundus View | 0 | 2 | 2 | 1 | 0 |
|  | **Intravitreal** | Anterior Chamber Flare | 0 | 0 | 0 | 0 | 0 |
|  |  | Anterior Chamber Cell | 0 | 0 | 0 | 0 | 0 |
|  |  | Anterior Vitreous Cell | 0 | 3 (Mixed) | 3 (Mixed) | 3 (Brown) | 1 (Brown) |
|  |  | Degraded Fundus View | 0 | 2 | 1 | 1 | 0 |
